## Supplementary Material for "Fragment ion intensity prediction improves the identification rate of non-tryptic peptides in timsTOF"

### Supplementary Figures

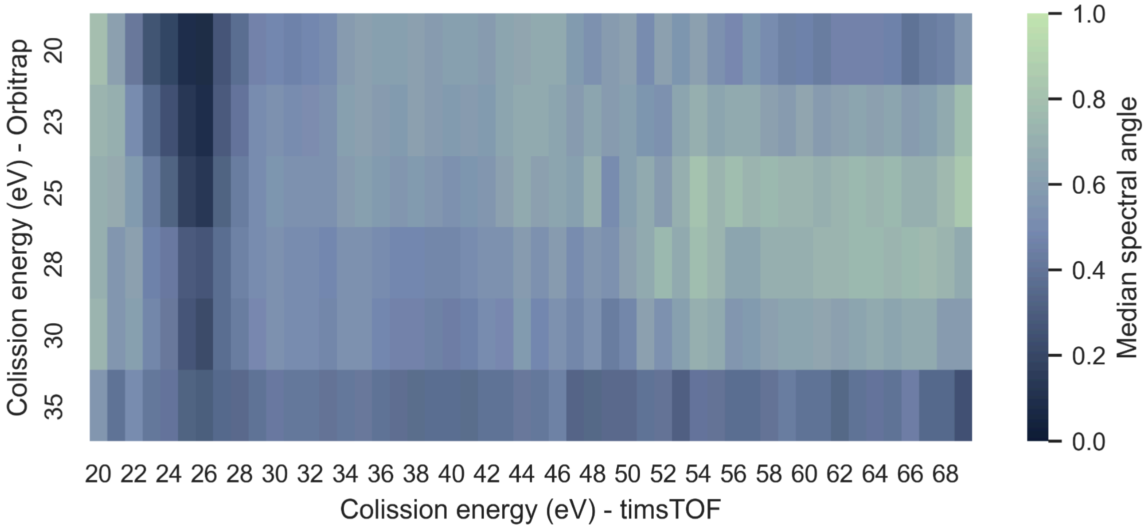


**Supplementary Figure S1.** The dissimilarity between MS/MS spectra generated by timsTOF and Orbitrap instruments for the same peptides was assessed by calculating the normalized spectral contrast angle between all MS/MS spectra with the same peptide sequence and charge state. The Orbitrap spectra originated from the hold-out set of Wilhelm et al. [1]. The collision energies do not seem to translate between the two machines.

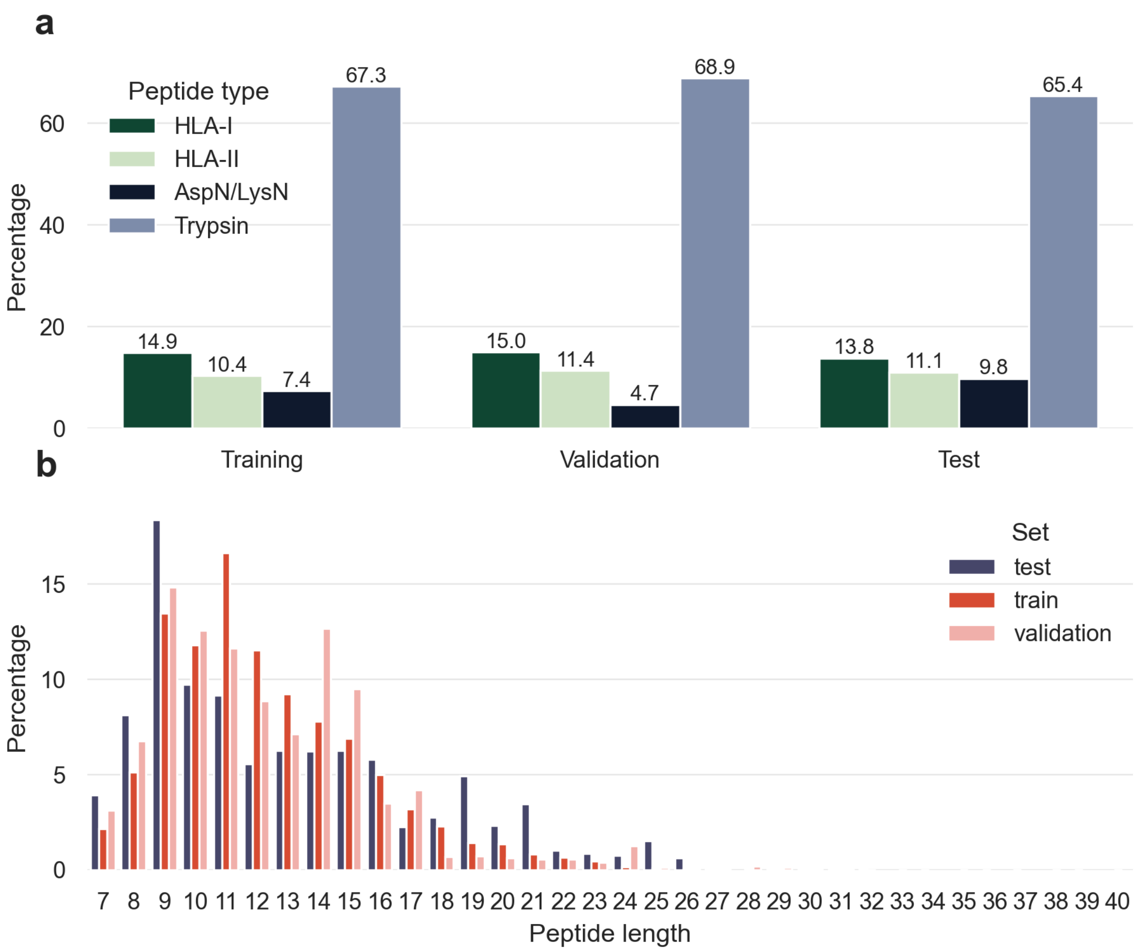


**Supplementary Figure S2.** Descriptive analysis of the compiled data. **a** Distribution of the different peptide types across the training, validation, and test sets. **b** Distribution of the peptide lengths across the training, validation, and test sets.

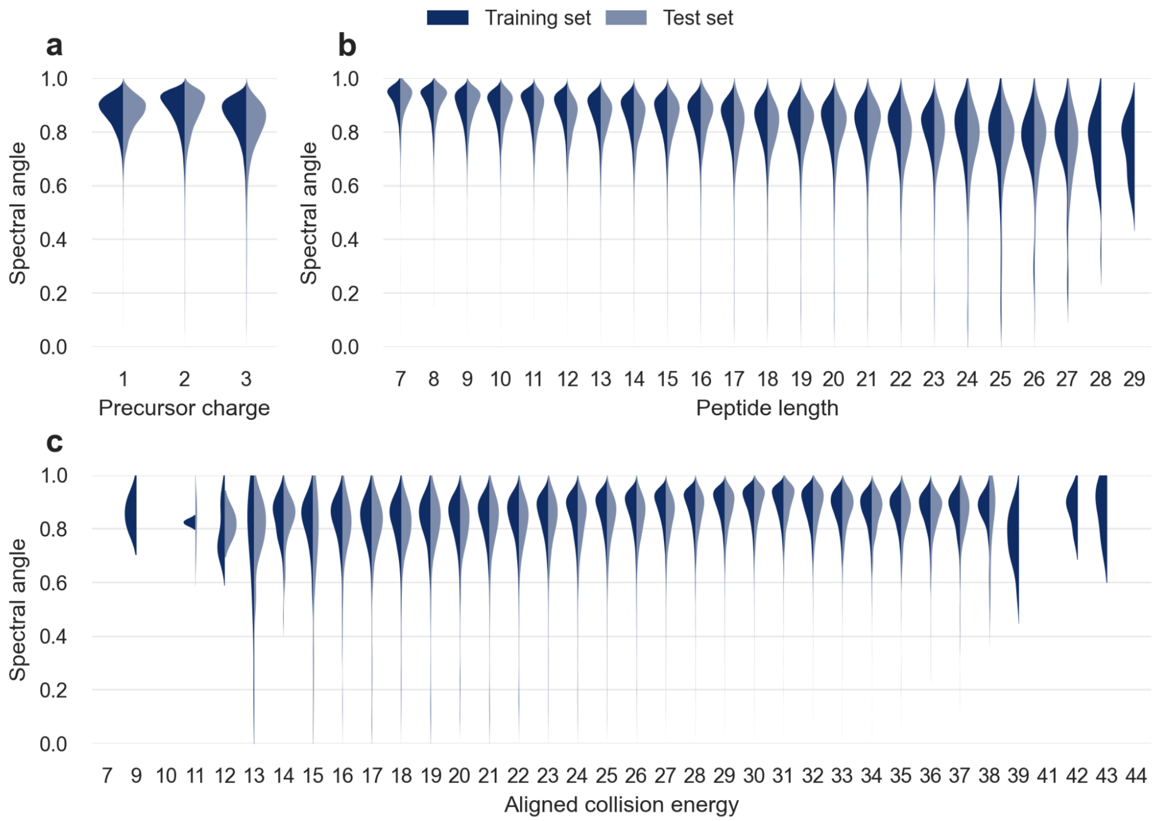


**Supplementary Figure S3.** The performance of the timsTOF Prosit 2023 model was assessed using asymmetric violin plots depicting the normalized spectral contrast angle between predicted and observed spectra in both the training and test sets. The model performed well across different precursor charges (**a**) and a wide range of collision energies (**c**) with only a minor influence of these factors on the predicted fragment ion intensities (Pearson correlation coefficients of -0.19 and 0.22 for the precursor charge and the collision energy, respectively). The model did show a moderate influence of the peptide length (**b**) on the predicted fragment ion intensities (Pearson correlation coefficient of -0.38).

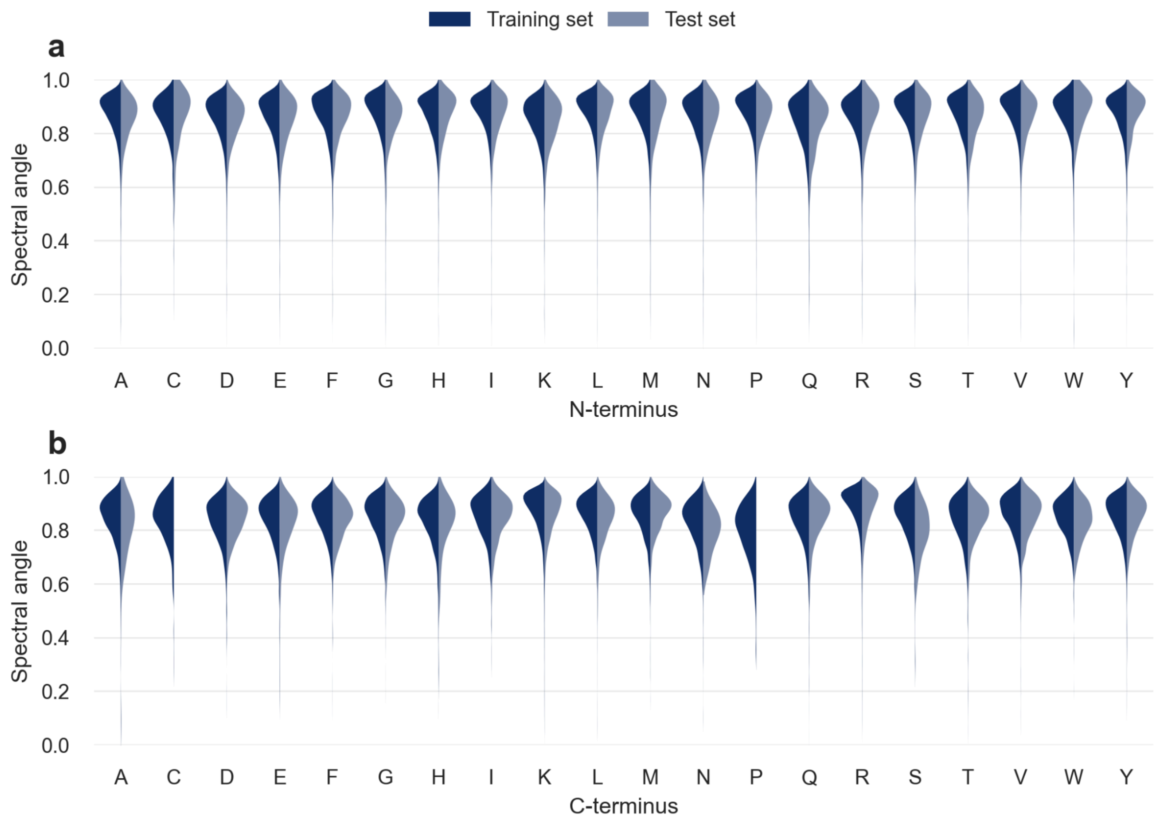


**Supplementary Figure S4.** The timsTOF Prosit 2023 model showed little bias for (**a)** N- and, (**b**) C-terminal amino acids.

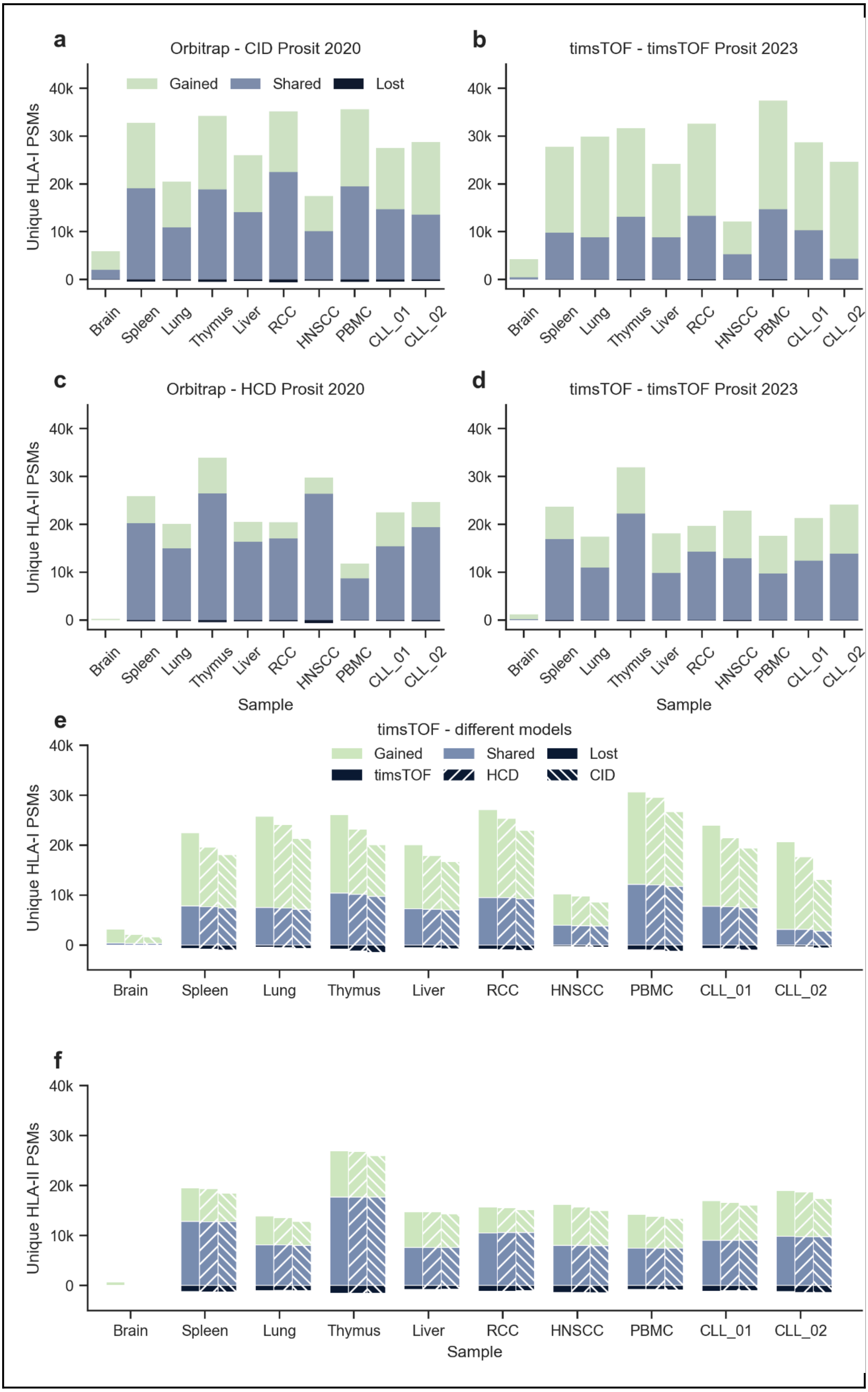
a
Orbitrap - CID Prosit 2020
Gained
Shared
Lost
Unique HLA-I PSMs
40k
30k
20k
10k
0
Brain
Spleen
Lung
Thymus
Liver
RCC
HNSCC
PBMC
CLL_01
CLL_02
b
timsTOF - timsTOF Prosit 2023
c
Orbitrap - HCD Prosit 2020
Unique HLA-II PSMs
Sample
d
timsTOF - timsTOF Prosit 2023
Sample
e
timsTOF - different models
Gained
Shared
Lost
timsTOF
HCD
CID
Unique HLA-I PSMs
f
Unique HLA-II PSMs
Sample

**Supplementary Figure S5. Gained, shared, and lost identified PSMs for different sample types to compare PSM rescoring on Orbitrap data with PSM rescoring on timsTOF data.** In general PSM rescoring was able to boost the number of PSMs, retaining true PSMs, gaining new PSMs, and losing only a small number of previously incorrect PSMs. **a** On average PSM rescoring of HLA-I Orbitrap data with the CID Prosit 2020 model resulted in a 1.9-fold increase (with 145,280 shared, 118,453 gained, and 3,466 lost PSMs). **b** On average PSM rescoring of HLA-I timsTOF data with the timsTOF Prosit 2023 model resulted in a 3.6-fold increase (with 89,098 shared, 163,852 gained, and 863 lost PSMs). **c** On average PSM rescoring of HLA-II Orbitrap data with the HCD Prosit 2020 model resulted in a 1.3-fold increase (with 165,160 shared, 44,796 gained, and 2,749 lost PSMs). **d** On average PSM rescoring of HLA-II timsTOF data with the timsTOF Prosit 2023 model resulted in a 2.3-fold increase (with 123,232 shared, 74,587 gained, and 1,328 lost PSMs). **e** To evaluate the effect of the fragment ion intensity prediction model on PSM rescoring, the RT prediction-based features were excluded. On average PSM rescoring of HLA-I timsTOF data with the timsTOF Prosit 2023 model resulted in a 3.4-fold increase (with 69,832 shared, 140,444 gained, and 5,437 lost PSMs). On average PSM rescoring of HLA-I timsTOF data with the HCD Prosit 2020 model resulted in a 2.9-fold increase (with 68,802 shared, 122,033 gained, and 6,467 lost PSMs). On average PSM rescoring of HLA-I timsTOF data with the CID Prosit 2020 model resulted in a 2.5-fold increase (with 66,745 shared, 101,782 gained, and 8,524 lost PSMs). **f** To evaluate the effect of the fragment ion intensity prediction model on PSM rescoring, the RT prediction-based features were excluded. On average PSM rescoring of HLA-II timsTOF data with the timsTOF Prosit 2023 model resulted in a 1.6-fold increase (with 91,281 shared, 66,874 gained, and 10,256 lost PSMs). On average PSM rescoring of HLA-II timsTOF data with the HCD Prosit 2020 model resulted in a 1.6-fold increase (with 91,184 shared, 63,897 gained, and 10,353 lost PSMs). On average PSM rescoring of HLA-II timsTOF data with the CID Prosit 2020 model resulted in a 1.5-fold increase (with 91,111 shared, 57,688 gained, and 10,426 lost PSMs). RCC = renal cell carcinoma; HNSCC = head and neck squamous-cell carcinoma; PBMC = peripheral blood mononuclear cell; CLL = chronic lymphocytic leukemia.

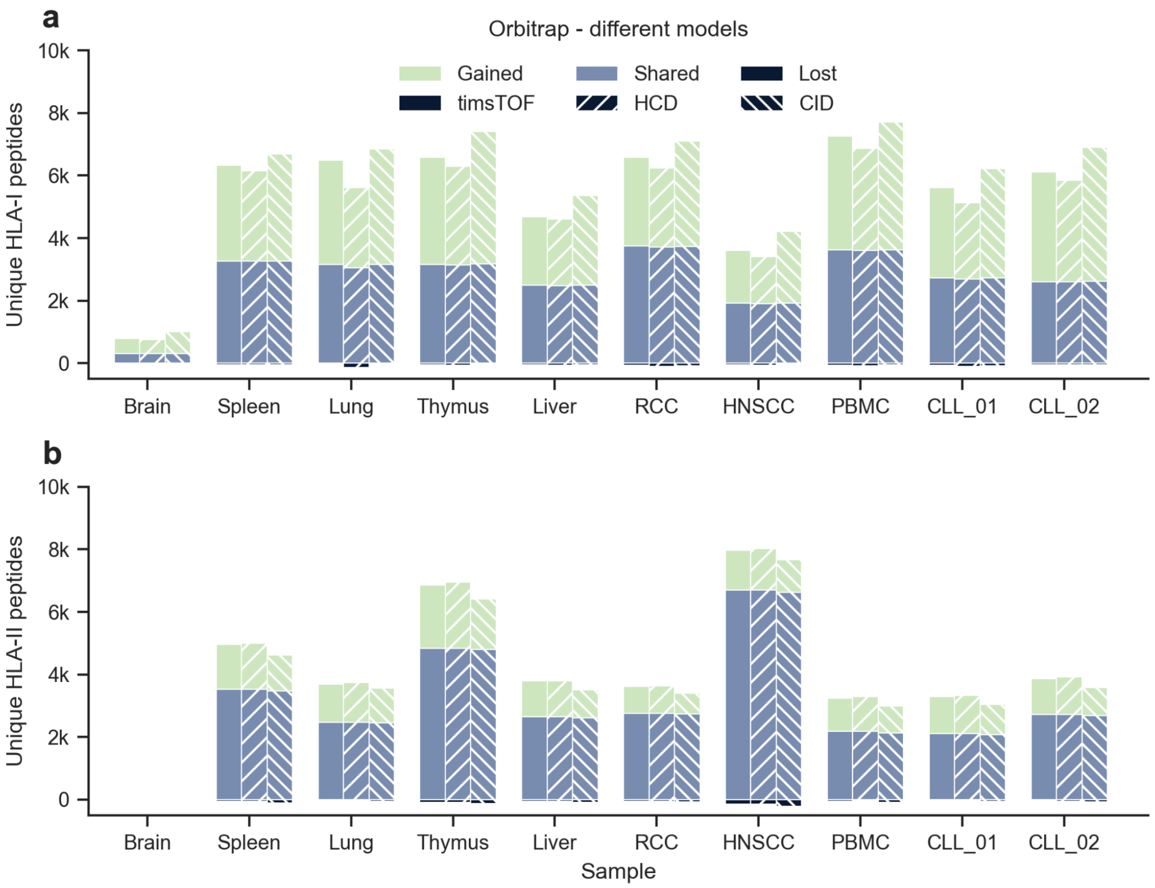


**Supplementary Figure S6. Gained, shared, and lost peptide identifications for PSM rescoring with different fragment ion intensity prediction models.** In general PSM rescoring was able to boost the number of PSMs, retaining true PSMs, gaining new PSMs, and losing only a small number of previously incorrect PSMs. **a** To evaluate the effect of the fragment ion intensity prediction model on PSM rescoring, the RT prediction-based features were excluded. On average PSM rescoring of HLA-I Orbitrap data with the timsTOF Prosit 2023 model resulted in a 2.0-fold increase (with 26,994 shared, 27,056 gained, and 511 lost PSMs). On average PSM rescoring of HLA-I timsTOF data with the HCD Prosit 2020 model resulted in a 1.9-fold increase (with 26,775 shared, 24,137 gained, and 730 lost PSMs). On average PSM rescoring of HLA-I timsTOF data with the CID Prosit 2020 model resulted in a 2.3-fold increase (with 27,009 shared, 32,462 gained, and 496 lost PSMs). **b** To evaluate the effect of the fragment ion intensity prediction model on PSM rescoring, the RT prediction-based features were excluded. On average PSM rescoring of HLA-II timsTOF data with the timsTOF Prosit 2023 model resulted in a 1.4-fold increase (with 29,945 shared, 11,305 gained, and 569 lost PSMs). On average PSM rescoring of HLA-II timsTOF data with the HCD Prosit 2020 model resulted in a 1.4-fold increase (with 29,947 shared, 11,742 gained, and 567 lost PSMs). On average PSM rescoring of HLA-II timsTOF data with the CID Prosit 2020 model resulted in a 1.3-fold increase (with 29,605 shared, 9,162 gained, and 909 lost PSMs). RCC = renal cell carcinoma; HNSCC = head and neck squamous-cell carcinoma; PBMC = peripheral blood mononuclear cell; CLL = chronic lymphocytic leukemia.

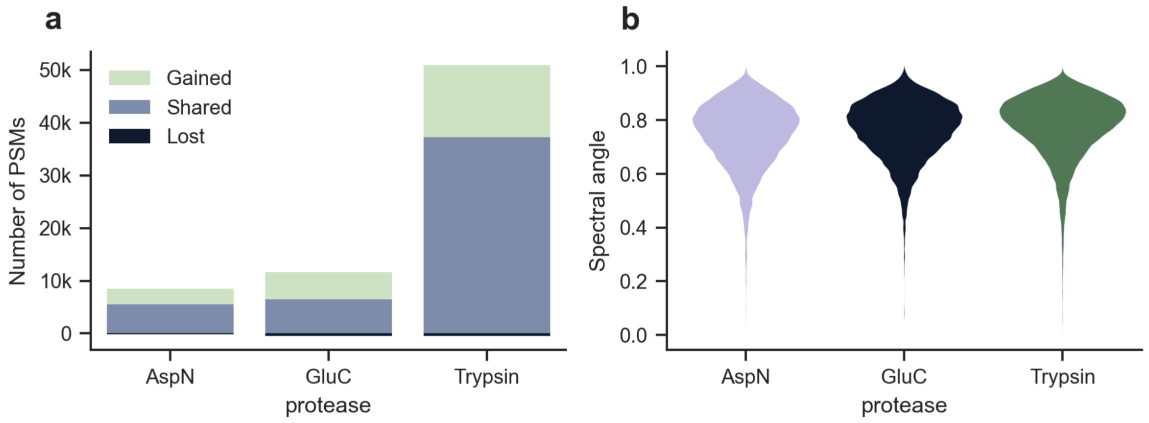


**Supplementary figure S7. PSM rescoring of samples cleaved with either AspN, GluC, or trypsin.** In general PSM rescoring was able to boost the annotation rate for all three samples, retaining true PSMs, gaining new PSMs, and losing only a small number of previously incorrect PSMs. **a** PSM rescoring of the sample cleaved with AspN resulted in a 1.5-fold increase (with 2,983 gained, 5,512 shared, and 176 lost PSMs). PSM rescoring of the sample cleaved with GluC resulted in a 1.7-fold increase (with 5,189 gained, 6,454 shared, and 496 lost PSMs). PSM rescoring of the sample cleaved with trypsin resulted in a 1.4-fold increase (with 13,624 gained, 37,253 shared, and 489 lost PSMs). **b** The performance of the timsTOF Prosit 2023 model was assessed using violin plots depicting the normalized spectral contrast angle between predicted and observed spectra. The median spectral angle observed for AspN, GluC, and trypsin respectively, was 0.77, 0.78, and 0.80.

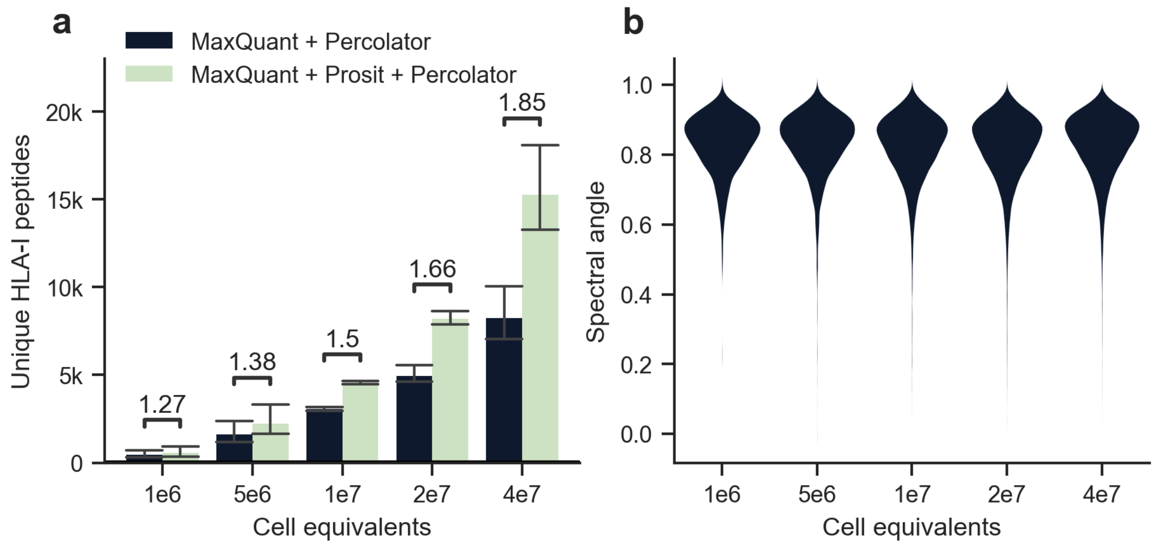


**Supplementary Figure S8. a** Mean and standard deviation showing the number of unique HLA-I peptides identified from 1 million to 40 million A-375 cell equivalents. Above each bar the fold change is shown between MaxQuant + Percolator and MaxQuant + Prosit + Percolator. **b** Violin plots depicting the normalized spectral contrast angle between predicted and observed spectra. The median spectral angle observed for cell equivalents from 1 million to 40 million, respectively, were 0.85, 0.85, 0.84, 0.84, and 0.85.

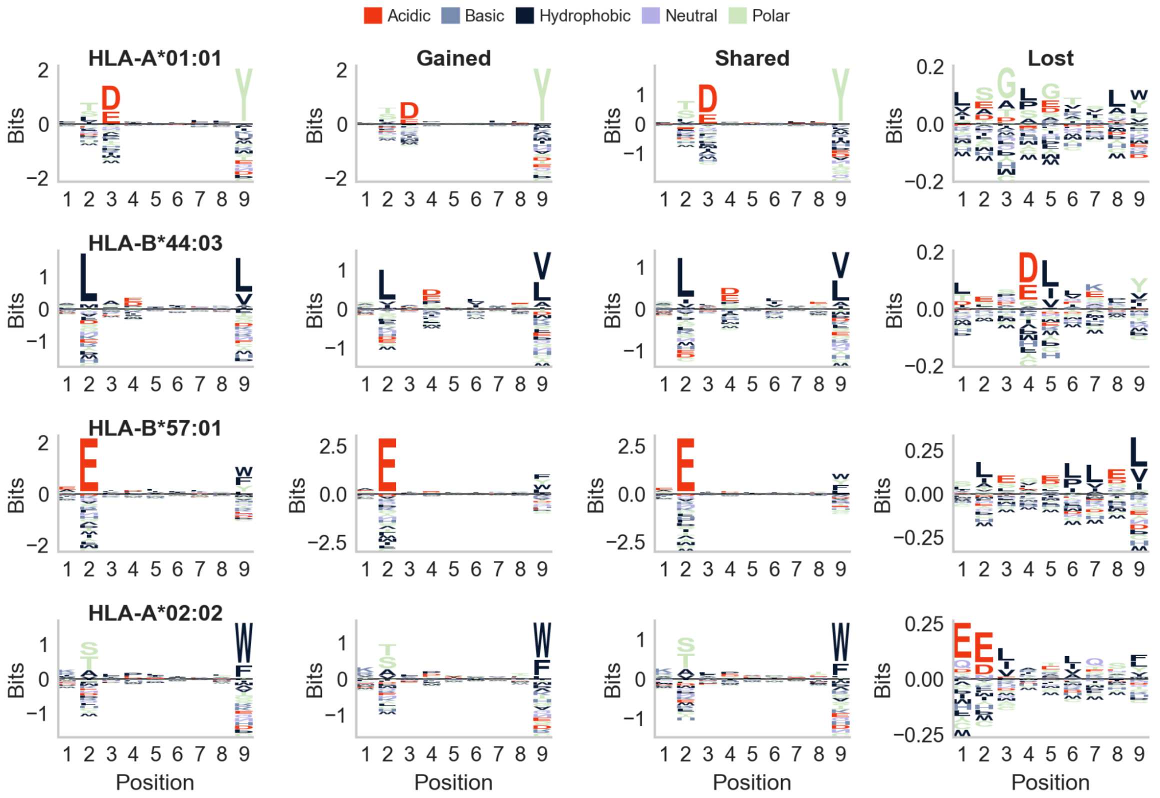


**Supplementary Figure S9.** HLA motif plots were generated for four monoallelic cell lines expressing specific HLA alleles: A*01:01 (1,406 unique peptides), A*02:02 (3,483 unique peptides), B*44:03 (989 unique peptides), and B*57:01 (1,510 unique peptides) [2]. From the PSM rescoring results of the 1 million to 40 million cell equivalents, a total of 16,641 unique gained peptides, 13,208 unique shared peptides, and 447 unique lost peptides were clustered, resulting in four peptide motifs for each peptide set. Next to each HLA motif, the peptide motif is plotted with the smallest Kullback-Leibler distance compared to the HLA motif. Amino acids are colored according to their physico-chemical properties (red acidic, blue basic, black hydrophobic, purple neutral, and green polar amino acids).

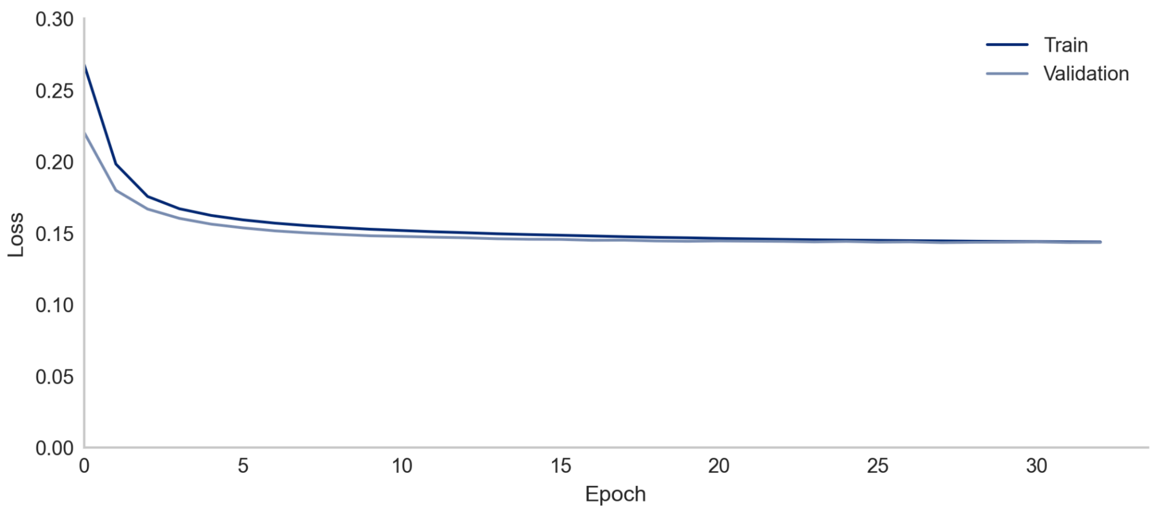


**Supplementary Figure S8. Loss plot illustrating the model training progress.** The loss function was 1 minus the normalized spectral contrast angle. The plot displays the train and validation loss values across the epochs, offering insights into the convergence and performance of model training over time.

| **Feature** | **Description** |
| --- | --- |
| SpecId | Internal spectrum ID |
| Label | 1 indicates target, -1 decoy |
| ScanNr | Scan number hash |
| filename | Raw file name |
| ExpMass | Fixed constant to allow PSMs with different theoretical mass to compete for the same PSM in target-decoy competition |
| CID | 1 indicates CID fragmentation |
| Charge1 | Boolean for charge state 1 |
| Charge2 | Boolean for charge state 2 |
| Charge3 | Boolean for charge state 3 |
| Charge4 | Boolean for charge state 4 |
| Charge5 | Boolean for charge state 5 |
| Charge6 | Boolean for charge state 6 |
| HCD | 1 indicates HCD fragmentation |

| | |
|---|---|
| KR | Number of K or R amino acids in sequence |
| Mass | Experimental mass |
| UnknownFragmentationMethod | 1 indicates unknown fragmentation method |
| andromeda | Score returned by the search engine |
| andromeda_delta_score | Delta score from the best next peptide identification returned by the search engine |
| missedCleavages | Number of missed cleavages |
| sequence_length | Peptide sequence length |
| Peptide | Peptide sequence |
| Protein | Protein description |

**Supplementary Table S2. List of features used for PSM rescoring without Prosit predictions.** Both the Features and the descriptions are reported. These features were used to postprocess the MaxQuant results to have a fair comparison with the

| **Feature** | **Description** |
|---|---|
| SpecId | Internal spectrum ID |
| Label | 1 indicates target, -1 decoy |
| ScanNr | Scan number |
| filename | Raw file name |
| ExpMass | Fixed constant to allow PSMs with different theoretical mass to compete for the same PSM in target-decoy competition |
| CID | 1 indicates CID fragmentation |
| Charge1 | Boolean for charge state 1 |
| Charge2 | Boolean for charge state 2 |
| Charge3 | Boolean for charge state 3 |
| Charge4 | Boolean for charge state 4 |
| Charge5 | Boolean for charge state 5 |
| Charge6 | Boolean for charge state 6 |
| HCD | 1 indicates HCD fragmentation |

| | |
|---|---|
| KR | Number of K or R amino acids in sequence |
| Mass | Experimental mass |
| RT | Experimental retention time |
| UnknownFragmentationMethod | 1 indicates unknown fragmentation method |
| abs_diff_Q1 | Quantile 1 absolute differences between the predicted and theoretical ions |
| abs_diff_Q2 | Quantile 2 absolute differences between the predicted and theoretical ions |
| abs_diff_Q3 | Quantile 3 of the absolute differences between the predicted and theoretical ions |
| collision_energy_aligned | Selected aligned collision energy |
| cos | Cosine similarity on all potential y- and b-ions |
| count_not_observed_and_not_predicted | Number of theoretical ions not observed in the experimental spectrum nor in the predicted spectrum |
| count_not_observed_and_not_predicted_b | Number of theoretical b-ions not observed in the experimental spectrum nor found in the predicted spectrum |
| count_not_observed_and_not_predicted_y | Number of theoretical y-ions not observed in the experimental spectrum nor found in the predicted spectrum |
| count_not_observed_but_predicted | Number of theoretical ions not observed in the experimental spectrum but found in the predicted spectrum |
| count_not_observed_but_predicted_b | Number of theoretical b-ions not observed in the experimental spectrum but found in the predicted spectrum |
| count_not_observed_but_predicted_y | Number of theoretical y-ions not observed in the experimental spectrum but found in the predicted spectrum |
| count_observed | Number observed annotated ions in the experimental spectrum |
| count_observed_and_predicted | Number observed annotated ions in the experimental spectrum and found in the predicted spectrum |

| | |
|---|---|
| count_observed_and_predicted_b | Number observed annotated b-ions in the experimental spectrum and found in the predicted spectrum |
| count_observed_and_predicted_y | Number observed annotated y-ions in the experimental spectrum and found in the predicted spectrum |
| count_observed_b | Number observed annotated b-ions in the experimental spectrum |
| count_observed_but_not_predicted | Number observed annotated ions in the experimental spectrum and not found in the predicted spectrum |
| count_observed_but_not_predicted_b | Number observed annotated b-ions in the experimental spectrum and not found in the predicted spectrum |
| count_observed_but_not_predicted_y | Number observed annotated y-ions in the experimental spectrum and not found in the predicted spectrum |
| count_observed_y | Number observed annotated y-ions in the experimental spectrum |
| count_predicted | Number of the predicted ions |
| count_predicted_b | Number of the predicted b-ions |
| count_predicted_y | Number of the predicted y-ions |
| fraction_not_observed_and_not_predicted | Number of theoretical ions not observed in the experimental spectrum nor found in the predicted spectrum divided by the number of theoretical ions |
| fraction_not_observed_and_not_predicted_b | Number of theoretical b-ions not observed in the experimental spectrum nor found in the predicted spectrum divided by the number of theoretical b-ions |
| fraction_not_observed_and_not_predicted_b_vs_predicted_b | Number of theoretical b-ions not observed in the experimental spectrum nor found in the predicted spectrum divided by the number of b-ions found in the predicted spectrum |
| fraction_not_observed_and_not_predicted_vs_predicted | Number of theoretical ions not observed in the experimental spectrum nor found in the predicted spectrum divided by the number of ions found in the predicted spectrum |
| fraction_not_observed_and_not_predicted_y | Number of theoretical y-ions not observed in the experimental spectrum nor found in the predicted |

| | |
|---|---|
| | spectrum divided by the number of theoretical y-ions |
| fraction_not_observed_and_not_predicted_y_vs_predicted_y | Number of theoretical y-ions not observed in the experimental spectrum nor found in the predicted spectrum divided by the number of y-ions found in the predicted spectrum |
| fraction_not_observed_but_predicted | Number of theoretical ions not observed in the experimental spectrum but found in the predicted spectrum divided by the number of theoretical ions |
| fraction_not_observed_but_predicted_b | Number of theoretical b-ions not observed in the experimental spectrum but found in the predicted spectrum divided by the number of theoretical b-ions |
| fraction_not_observed_but_predicted_b_vs_predicted | Number of theoretical b-ions not observed in the experimental spectrum but found in the predicted spectrum divided by the number of b-ions found in the predicted spectrum |
| fraction_not_observed_but_predicted_vs_predicted | Number of theoretical ions not observed in the experimental spectrum but found in the predicted spectrum divided by the number of ions found in the predicted spectrum |
| fraction_not_observed_but_predicted_y | Number of theoretical y-ions not observed in the experimental spectrum but found in the predicted spectrum divided by the number of theoretical y-ions |
| fraction_not_observed_but_predicted_y_vs_predicted | Number of theoretical y-ions not observed in the experimental spectrum but found in the predicted spectrum divided by the number of y-ions found in the predicted spectrum |
| fraction_observed | Number of theoretical ions observed in the experimental spectrum divided by the number of theoretical ions |
| fraction_observed_and_predicted | Number of theoretical ions observed in the experimental spectrum and found in the predicted spectrum divided by the number of theoretical ions |
| fraction_observed_and_predicted_b | Number of theoretical b-ions observed in the experimental spectrum and found in the predicted spectrum divided by the number of theoretical b-ions |

| | |
|---|---|
| fraction_observed_and_predicted_b_vs_predicted_b | Number of theoretical b-ions observed in the experimental spectrum and found in the predicted spectrum divided by the number of predicted b-ions |
| fraction_observed_and_predicted_vs_predicted | Number of theoretical ions observed in the experimental spectrum and found in the predicted spectrum divided by the number of predicted ions |
| fraction_observed_and_predicted_y | Number of theoretical y-ions observed in the experimental spectrum and found in the predicted spectrum divided by the number of theoretical y-ions |
| fraction_observed_and_predicted_y_vs_predicted_y | Number of theoretical y-ions observed in the experimental spectrum and found in the predicted spectrum divided by the number of predicted y-ions |
| fraction_observed_b | Number of theoretical b-ions observed in the experimental spectrum divided by the number of theoretical b-ions |
| fraction_observed_but_not_predicted | Number of theoretical ions observed in the experimental spectrum but not found in the predicted spectrum divided by the number of theoretical ions |
| fraction_observed_but_not_predicted_b | Number of theoretical b-ions observed in the experimental spectrum but not found in the predicted spectrum divided by the number of theoretical b-ions |
| fraction_observed_but_not_predicted_b_vs_predicted_b | Number of theoretical b-ions observed in the experimental spectrum but not found in the predicted spectrum divided by the number of predicted b-ions |
| fraction_observed_but_not_predicted_vs_predicted | Number of theoretical ions observed in the experimental spectrum but not found in the predicted spectrum divided by the number of predicted ions |
| fraction_observed_but_not_predicted_y | Number of theoretical y-ions observed in the experimental spectrum but not found in the predicted spectrum divided by the number of theoretical y-ions |
| fraction_observed_but_not_predicted_y_vs_predicted_y | Number of theoretical y-ions observed in the experimental spectrum but not found in the predicted spectrum divided by the number of predicted y-ions |

| | |
|---|---|
| fraction_observed_y | Number of theoretical y-ions observed in the experimental spectrum divided by the number of theoretical y-ions |
| fraction_predicted | Number of theoretical ions found in the predicted spectrum divided by the number of theoretical ions |
| fraction_predicted_b | Number of theoretical b-ions found in the predicted spectrum divided by the number of theoretical b-ions |
| fraction_predicted_y | Number of theoretical y-ions found in the predicted spectrum divided by the number of theoretical y-ions |
| max_abs_diff | Max absolute difference between the predicted and theoretical ions |
| mean_abs_diff | Mean absolute difference between the predicted and theoretical ions |
| min_abs_diff | Minimum absolute difference between the predicted and theoretical ions |
| missedCleavages | Number of missed cleavages |
| modified_cosine | Modified cosine similarity [3] on all potential y- and b-ions |
| mse | Mean square error |
| pearson_corr | Pearson correlation on all potential y- and b-ions |
| pearson_corr_b_ions | Pearson correlation on b-ions |
| pearson_corr_double_charge | Pearson correlation on doubly charged ions |
| pearson_corr_single_charge | Pearson correlation on singly charged ions |
| pearson_corr_triple_charge | Pearson correlation on triply charged ions |
| pearson_corr_y_ions | Pearson correlation on y-ions |
| sequence_length | Peptide sequence length |
| spearman_corr | Spearman correlation on all potential y- and b-ions |
| spearman_corr_b_ions | Spearman correlation on b-ions |
| spearman_corr_double_charge | Spearman correlation on doubly charged ions |
| spearman_corr_single_charge | Spearman correlation on singly charged ions |

| | |
|---|---|
| spearman_corr_triple_charge | Spearman correlation on triply charged ions |
| spearman_corr_y_ions | Spearman correlation on y-ions |
| spectral_angle | Normalized spectral contrast angle (SA) on all potential y- and b-ions |
| spectral_angle_b_ions | Normalized spectral contrast angle (SA) on b-ions |
| spectral_angle_double_charge | Normalized spectral contrast angle (SA) on doubly charged ions |
| spectral_angle_single_charge | Normalized spectral contrast angle (SA) on singly charged ions |
| spectral_angle_triple_charge | Normalized spectral contrast angle (SA) on triply charged ions |
| spectral_angle_y_ions | Normalized spectral contrast angle (SA) on y-ions |
| spectral_entropy_similarity | Spectral entropy similarity [4] on all potential y- and b-ions |
| std_abs_diff | Standard deviation of the absolute differences between retention time and aligned predicted retention time |
| abs_rt_diff | Absolute difference between retention time and aligned predicted retention time |
| lda_scores | Score returned by a linear discriminant analysis on the spectral angle to estimate false discovery rates |
| pred_RT | Predicted aligned retention time |
| iRT | Predicted indexed retention time |
| Peptide | Peptide sequence |
| Protein | Protein description |

**Supplementary Table S1. List of features used for PSM rescoring with Prosit predictions.** Both the features and the descriptions are reported. When fragment ion intensity prediction models were compared, the following features were removed: abs_rt_diff, lda_scores, pred_RT, and iRT.

### References

1. Wilhelm, M. *et al.* Deep learning boosts sensitivity of mass spectrometry-based immunopeptidomics. *Nat. Commun.* **12**, 3346 (2021).
2. Sarkizova, S. *et al.* A large peptidome dataset improves HLA class I epitope prediction across most of the human population. *Nat. Biotechnol.* **38**, 199–209 (2020).
3. McGann, C. D. *et al.* Real-Time Spectral Library Matching for Sample Multiplexed Quantitative Proteomics. *J. Proteome Res.* **22**, 2836–2846 (2023).
4. Li, Y. *et al.* Spectral entropy outperforms MS/MS dot product similarity for small-molecule compound identification. *Nat. Methods* **18**, 1524–1531 (2021).
